# Landscape genetics across the Andes Mountains: Environmental variation drives genetic divergence in the leaf-cutting ant *Atta cephalotes*

**DOI:** 10.1101/2021.12.19.473407

**Authors:** Vanessa Muñoz-Valencia, James Montoya-Lerma, Perttu Seppä, Fernando Diaz

## Abstract

Disentangling the mechanisms underlying spatial distribution of genetic variation into those due to environment or physical barriers from mere geographic distance is challenging in complex landscapes. The Andean uplift represents one of the most heterogeneous habitats where these questions remain unexplored as multiple mechanisms might interact, confounding their relative roles. We explore this broad question in the leaf-cutting ant *Atta cephalotes*, a species distributed across the Andes Mountains using nuclear microsatellite markers and *mtCOI* gene sequences. We investigate spatial genetic divergence across the Western range of Northern Andes in Colombia by testing the relative role of alternative scenarios of population divergence, including geographic distance (IBD), climatic conditions (IBE), and physical barriers due to the Andes Mountains (IBB). Our results reveal substantial genetic differentiation among *A. cephalotes* populations for both types of markers, but only nuclear divergence followed a hierarchical pattern with multiple models of genetic divergence imposed by the Western range. Model selection showed that IBD, IBE (temperature and precipitation), and IBB (Andes mountains) models, often proposed as individual drivers of genetic divergence, interact, and explain up to 33% of genetic divergence in *A. cephalotes.* IBE models remained significant after accounting for IBD, suggesting that environmental factors play a more prominent role when compared to IBB. These factors in combination with idiosyncratic dispersal patterns of ants appear to determine hierarchical patterns of gene flow. This study enriches our understanding of the forces shaping population divergence in complex habitat landscapes.

## INTRODUCTION

Population divergence can be determined by the spatial distribution of individuals where geographic proximity modulates genetic similarity, leading to a pattern of isolation by distance – IBD (Wright 1943; Slatkin 1993; Lee and Mitchell-Olds 2011; Shafer and Wolf 2013). However, in complex landscapes, populations are often exposed to environmental variation and physical barriers that may also contribute to genetic divergence (Manel et al. 2003; Manel and Holderegger 2013; Shafer and Wolf 2013). Populations might adapt to their local environment, maximizing the fitness of individuals under local conditions while decreasing the fitness of immigrants from alternative environments (Wang et al. 2013; Sobel 2014; Wang and Bradburd 2014; Carro et al. 2019). This adaptive reduction in gene flow can result in a pattern of isolation by environment – IBE (Shafer and Wolf 2013; Sexton et al. 2014; Wang and Bradburd 2014). Alternatively, gene flow might be restricted by allopatric scenarios of genetic differentiation mediated by physical barriers in the landscape, generating patterns of isolation by barrier – IBB (Haffer 2008; Rull 2011; Turchetto-Zolet et al. 2013; De Queiroz et al. 2017). When the complexity of landscapes increases, gene flow is likely influenced by a combination of geographical and ecological factors where these isolating mechanisms are not mutually exclusive (Crispo et al. 2006; Edwards et al. 2012; Wang et al. 2013; Noguerales et al. 2016).

The Andean Mountain ranges not only represent one of the most unexplored environments but also offer a great complexity of landscapes promoting diversification over a wide range of taxa (Salgado-Roa et al. 2018). Restricted gene flow across these mountains has been reported in multiple organisms, such as birds (Cadena et al. 2016), plants (Luebert and Weigend 2014; Lagomarsino et al. 2016; Pérez-Escobar et al. 2017), mammals (Antonelli et al. 2009; Hoorn et al. 2010), insects (Antonelli et al. 2009; Hoorn et al. 2010; De-Silva et al. 2017) and other arthropods (Salgado-Roa et al. 2018). However, organisms form contrasting populations in these habitats are often exposed to a combination of distance, environment, and physical barriers to dispersal, challenging the investigation on the relative role of the different isolating mechanisms (James et al. 2011; Meirmans 2015; Noguerales et al. 2016). Therefore, it is still unclear how the Andean uplift has modulated patterns of gene flow and the evolution of several of the most ecologically important groups in the Neotropics, including social insects. For instance, although the entire evolution of Neotropical ants takes place across the Andes (Mueller et al. 2017), the interplay between their population structure and environmental variation relative to the effect of these mountains in isolated populations, remains largely unknown.

The leaf-cutting ant *A. cephalotes* is a major urban and agricultural pest in the Neotropics, colonizing a wide spectrum of environments (Hölldobler and Wilson 2011; Della Lucia et al. 2014; Fernández et al. 2015). In Colombia, its distribution overlaps with the maximum complexity of the Andean uplift, ranging from the sea level up to 2,100 m.a.s.l. (Fernández et al. 2015), with a vertical thermal gradient of 0.6 °C/100 m (Hermelin 2015). The northern section of these mountains in Colombia splits into three main branches: the Western range, the Central range, and the Eastern range. *A. cephalotes’* populations are separated by these mountains while being exposed to complex combinations of topographical and environmental variation (Kattan et al. 2004; Pérez-Escobar et al. 2017; Salgado-Roa et al. 2018). Such conditions provide a tremendous climatic spectrum for local adaptation, with an interplay between population dynamics and species-specific dispersal patterns (Hakala et al. 2019). The evolution under such environmental heterogeneity might shape patterns gene flow (Lee and Mitchell-Olds 2011; Wang et al. 2013; Noguerales et al. 2016; De Queiroz et al. 2017), which often results in more complex scenarios than genetic divergence merely due to IBD (Wright 1943; Slatkin 1993). For example, we recently found that the Eastern range of the Andes in Colombia plays a major role as a geographic barrier to historical gene flow, restricting *A. cephalotes’* dispersion from north to south (Muñoz-Valencia et al. 2021). Although this initial study demonstrates a significant influence of the Andes on population divergence by the leaf-cutting ant at a phylogeographic scale, the role of local adaptation occurring at more regional scales across the Andes remains untested.

This work focuses on a finer and more complex geographic scale of *A. cephalotes*’ distribution: The Western range of the Andean uplift in Colombia. We use a landscape genetic approach to investigate the role of geographic features and environmental variation in defining patterns of spatial genetic structure in *A. cephalotes*. Using nuclear (microsatellites) and mitochondrial (*mtCOI*) markers, we test the relative roles played by geographic distance, climate variation, and a major dispersal barrier (Western range) in modulating patterns of gene flow. As a monogynous species (single-queen), *A. cephalotes* presumably experiences long-distance nuptial flights that can potentially overcome isolating barriers (Moser 1967; Cherrett 1968; Helms 2018). Our results demonstrate that gene flow is limited by a complex interaction of the three isolating mechanisms (IBD, IBE, and IBB) rather than merely IBD, while IBE appears to play a primary role over IBB. Investigating the spatial genetic structure of a species facing an exceptionally heterogeneous environment helps to understand the evolution and diversity of this ecologically dominant group of ants in the Neotropics.

## MATERIALS AND METHODS

### Sampling

Ant sampling was carried out in the Colombian Pacific and Andean regions, which are separated by the Western Mountain range of Colombian Andes (Figure 1). The Pacific region is classified as a tropical rainforest with an extremely humid climate and an annual average temperature of 27°C. The Andean region was further divided into two groups: Andean 1 (800 – 1,050 m.a.s.l.) and Andean 2 (1,300 – 2,200 m.a.s.l.), with highly variable climate conditions. The inner valleys in Andean 1 are climatically classified as tropical savanna and tend to be dry (annual temperature: 25°C), while the range summits in Andean 2 are more humid, with a temperate climate and tropical monsoons (annual temperature: 21°C; Supplementary table ST1) (Köppen 1884; Hernández-Camacho 1992; Kattan et al. 2004; Peel et al. 2007; Chen and Chen 2013).

**Figure 1.**
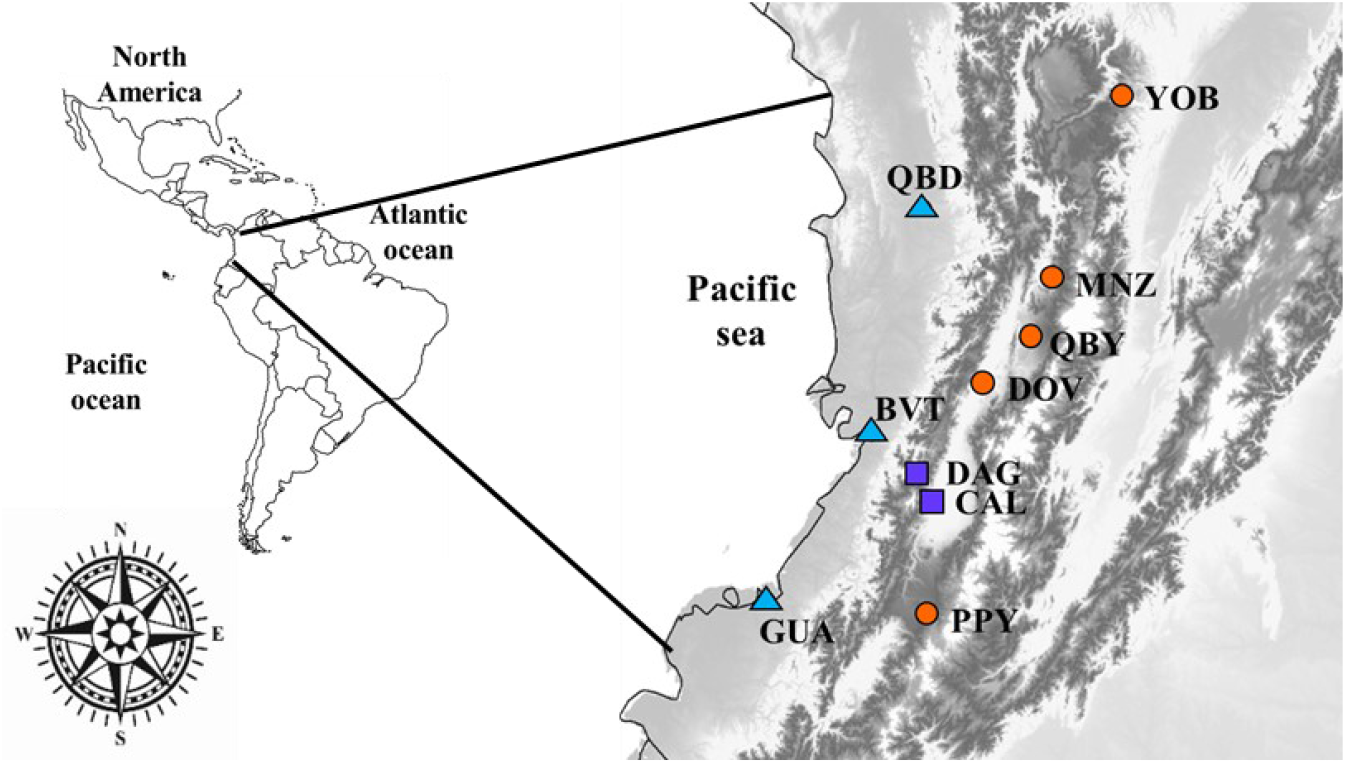
Study area and sampling locations of *Atta cephalotes* in Colombia. Different symbols and colors indicate sampling locations following regional classification: Blue triangles for sampling locations in the Pacific region (BVT, QBD, and GUA), violet squares for the Andean 1 region (DAG and CAL), and orange circles for the Andean 2 region (PPY, DOV, QBY, YOB, MNZ) (Supplementary table ST1).

Environmental variation across the regions was characterized by differences in temperature, humidity, and precipitation measured as five-year averages of the annual temperature (°C), relative humidity (%), and precipitation (mm) respectively for each location (IDEAM, 2019). In addition, a climate classification was represented using four categories (1 to 4) of different climatic conditions mediated by the tropical Andes. Variation in topography was estimated by elevation above sea level. A dummy variable was used to evaluate the Andean uplift as a major geographic barrier to gene flow. We used 0 to codify populations from the west side of the Western range (Pacific region), and 1 for populations from the east side of these mountains (Supplementary table ST1).

Worker ants were sampled between 2017-2018 from nine to twenty nests in ten locations from a total of 153 nests (Table 1). The between-nest distance per location was at least 1.5 km, ensuring that sampled nests were independent colonies. Three, two, and five locations were sampled from the Pacific, Andean 1, and Andean 2 regions, respectively (Table 1).

**Table 1.** Sampled populations of *A. cephalotes*. Locations with their regional classifications (Pacific, Andean 1 and Andean 2 regions from Colombia) and number of nests are indicated for each population.

| Code | Location | Region | # of nests |
| --- | --- | --- | --- |
| BVT | Buenaventura | Pacific | 9 |
| QBD | Quibdó | Pacific | 15 |
| GUA | Guapi | Pacific | 20 |
| DAG | Dagua | Andean_1 | 10 |
| CAL | Cali | Andean_1 | 20 |
| PPY | Popayán | Andean_2 | 14 |
| DOV | Dovio | Andean_2 | 19 |
| QBY | Quimbaya | Andean_2 | 16 |
| YOB | Yolombó | Andean_2 | 19 |
| MNZ | Manizales | Andean_2 | 11 |

### Molecular methods

#### DNA extraction and PCR amplification of microsatellite markers

Total DNA extraction was carried out for five workers from each nest (total 765 workers) using TNES lysis buffer (Tris 50mM, NaCl 0.4M, EDTA 100mM, SDS 0.5%) pH 7.5 and chloroform: isoamyl alcohol (24:1) following Wasko et al. (2003) with minor modifications as described by Muñoz-Valencia et al. (2020).

Thirteen microsatellite loci developed for *A. cephalotes* (Muñoz-Valencia et al. 2020) were used (Supplementary Table ST2). PCR reactions were carried out following Muñoz-Valencia et al. (2020) in a 10 μL volume containing 10 ng of DNA, 1 X of Phusion Flash PCR Master Mix (Thermo Fisher Scientific), and 2 μM of each labeled primer. The thermal profile was: 98 ºC for 1 min, 34 cycles of 98 °C for 1 sec, annealing temperature for 15 secs, and 72 ºC for 20 secs, followed by a final extension at 72 ºC for 1 min. The amplified fluorescent fragments were visualized using an automated DNA sequencer ABI 3130 Genetic Analyzer (Applied Biosystems), and allele sizes were estimated using GeneMapper version 4.0 (Thermo Fisher Scientific).

#### Sequencing of mitochondrial COI gene

A 368 bp fragment of the mitochondrial cytochrome oxidase subunit 1 gene (*mtCOI*) was sequenced using one sample per nest, obtaining a total of 146 sequences after discarding failed amplification and sequencing (GenBank accession numbers: MW245066 – MW245211). PCR amplification was carried out using universal primers Ben and Jerry, following Kronauer et al. (2004) and Simon et al. (1994). PCR reactions were performed in 10 μl volume, involving 10 ng of DNA, 1X GoTaq® Master Mix (PROMEGA), and 2 μM of each primer. The thermal cycle profile was carried out at 94 ºC for 2 min followed by 30 cycles of 94 ºC for 1 min, 58 ºC for 1 min, and 72 ºC for 1 min, with a final extension at 72 ºC for 10 min. PCR products were confirmed by 1 % agarose gels. All PCR products were sequenced by Psomagen, Maryland, USA.

### Population genetics analyses

#### Genetic diversity of DNA microsatellites and mtCOI gene

To test for null alleles, large allele dropout, and scoring errors, we used Microchecker version 2.2.3 (VanOosterhout et al. 2004). Putative null alleles were detected in five loci (MAT2, MAT4, MAT10, MAT15, MAT28) of five populations (DAG, GUA, DOV, QBY, YOB; Table 2), which may bias estimates of population structure (Chapuis and Estoup 2007). We accounted for this bias by comparing analyses after excluding these loci with those of the entire data set. Our main results and conclusions remained regardless of the loci included in the analyses, and only results for the entire data set are here reported. As *A. cephalotes* nests conform family units and nestmates are highly related, only one individual per nest was used to evaluate Hardy-Weinberg (HWE) and linkage disequilibrium (LD). Tests were performed for each locus (HWE) and each locus pair (LD) for each population (Supplementary Table ST2) using GENEPOP (Rousset 2008). A Bonferroni correction was applied with a global significance of α=0.05 for multiple comparisons.

**Table 2.** Microsatellites and mitochondrial *mtCOI* marker variation for sampling locations of *A. cephalotes* from Pacific, Andean 1 and Andean 2 regions from Colombia. *N* is the sample size (number of nests). *N_a_*: mean number of alleles; *AR*: mean allelic richness; *H_e_*: mean expected heterozygosity; *F*_IS_. inbreeding coefficient; *HWE*: Hardy-Weinberg Equilibrium test; *h*: number of haplotypes; *H_d_*. haplotype diversity; π: nucleotide diversity. (***) *P* < 0.0001.

|  | *Nuclear data* |  |  |  |  | *Mitochondrial data* |  |  |  |
| --- | --- | --- | --- | --- | --- | --- | --- | --- | --- |
| | *N* | *N<sub>a</sub>* | *AR* | *H<sub>e</sub>* | *F<sub>IS</sub>* | *N* | *h* | % *H<sub>d</sub>* | % $\pi$ |
| *Pacific* |  |  |  |  |  |  |  |  |  |
| All pops n = 3 | 44 | 6.95 | 6.56 | 0.57 | 0.01 | 42 | 5 | 67 | 0.4 |
| BVT | 9 | 6.08 | 6.02 | 0.59 | -0.06 | 9 | 2 | 22 | 0.2 |
| QBD | 15 | 7.46 | 6.97 | 0.58 | 0.00 | 13 | 4 | 60 | 0.4 |
| GUA | 20 | 7.31 | 6.68 | 0.55 | 0.04 | 20 | 4 | 56 | 0.4 |
| *Andean_1* |  |  |  |  |  |  |  |  |  |
| All pops n = 2 | 30 | 4.62 | 4.38 | 0.49 | -0.01 | 29 | 6 | 82 | 0.5 |
| DAG | 10 | 3.92 | 3.85 | 0.46 | -0.01 | 10 | 3 | 51 | 0.4 |
| CAL | 20 | 5.31 | 4.91 | 0.51 | -0.01 | 19 | 4 | 76 | 0.3 |
| *Andean_2* |  |  |  |  |  |  |  |  |  |
| All pops n = 5 | 79 | 4.74 | 4.51 | 0.53 | -0.01 | 75 | 6 | 53 | 0.2 |
| PPY | 14 | 3.69 | 3.58 | 0.54 | -0.01 | 14 | 3 | 62 | 0.2 |
| DOV | 19 | 4.38 | 4.18 | 0.48 | -0.05 | 19 | 3 | 29 | 0.1 |
| QBY | 16 | 4.77 | 4.58 | 0.52 | 0.02 | 15 | 2 | 25 | 0.1 |
| YOB | 19 | 6.38 | 5.83 | 0.61 | -0.01 | 16 | 1 | 0 | 0.0 |
| MNZ | 11 | 4.46 | 4.37 | 0.50 | 0.02 | 11 | 3 | 35 | 0.3 |
| **Total n = 10** | **153** | **10.38** | **7.06** | **0.60** | **-0.01** | **146** | **10** | **66** | **0.3** |

Genetic diversity was characterized for each locus, region, and population as allele count (N_a_), allelic richness (A_R_), expected (H_e_), and observed (H_o_) heterozygosities, and the inbreeding coefficient (*F*_IS_), using the software GenAlex v. 6.5 (Peakall and Smouse 2012) and FSTAT v. 2.9.3.2 (Goudet, 1995; 2001) (Table 2, Supplementary tables ST2, ST3). Tests for significant differences from zero of diversity parameters were performed using the *‘aov’* function in R (Chambers et al. 1992). We used the software Bottleneck (Piry et al. 1999) to estimate recent changes in population sizes by implementing the IAM and SSM mutation models and using a Wilcoxon signed-rank test to evaluate statistical significances. We also inspected whether the allele frequency distribution was shifted from the expected L-shape. When necessary, significant values were adjusted for multiple comparisons through Bonferroni correction.

Genetic diversity of *mtCOI* data was estimated through the number of haplotypes (h), nucleotide diversity (π), and haplotype diversity (h_d_) using DnaSP 5.19 (Librado and Rozas 2009). Tajima’s D and Fu’s FS neutrality tests were performed to investigate signatures of recent population expansion (Ramos-Onsins and Rozas 2002) when the null hypothesis of neutrality was rejected due to significant negative values (*P* < 0.05 for D, *P* < 0.02 for FS; Supplementary table ST3).

#### Spatial genetic patterns

We investigated spatial genetic patterns and restrictions to gene flow in *A. cephalotes* using the software ARLEQUIN version 3.5.2.2 (Excoffier and Lischer 2010) to perform alternative hierarchical Analysis of Molecular Variance (AMOVA). First, populations were classified into two regions separated by the Western range (Pacific, Andean) to study the effect of the Western range of Colombian Andes as a major geographic barrier to gene flow affecting the distribution of genetic variation in the data (isolation by barrier, IBB). Second, populations were classified into three regions defined by their climatic conditions mediated by the Andes mountains [(Pacific, Andean 1, Andean 2) isolation by environment, IBE; Supplementary table ST1]. In the AMOVA, we estimated associated *F*_ST_ for microsatellites and genetic distance-based *Φ*-statistic for *mtCOI* (Excoffier et al. 1992; Meirmans 2006; Meirmans and Hedrick 2011), both globally and between all pairs of populations. The significance of variance components and associated *F*_ST_ and *Φ* indices were calculated using 10,000 permutations.

We further investigated the structure of the *A. cephalotes* nuclear data using two clustering analyses. First, we used a model-based Bayesian clustering method implemented in the software STRUCTURE v. 2.3.4 (Pritchard et al. 2000), which estimates the number of genetic clusters (*K*) independent of spatial sampling. Analyses were performed using one individual per nest, with and without admixture, for correlated allele frequencies. A 50,000 burn-in and 500,000 sampling generations were implemented for *K* ranging from 1 to 12, with 10 iterations for each value of *K.* The Evanno’s method (Evanno et al. 2005) implemented in STRUCTURE HARVESTER (Earl and vonHoldt 2012) was used to estimate the optimal number of clusters from the STRUCTURE output (Supplementary figure SF1) and results were visualized using CLUMPAK (Kopelman et al. 2015).

Second, discriminant analysis of principal components (DAPC) was performed by using a principal component analysis (PCA) before the discriminant analysis (DA) (Jombart et al. 2010). DA partitions genetic variation, maximizing differences between clusters while minimizing within-cluster variation. We performed a DAPC analysis in the R package ADEGENET (Jombart et al. 2010) using one individual per nest for microsatellite data and polymorphic nucleotide positions matrix for *mtCOI.* The function *“dapc”* was used to estimate all available principal components (PCs) as well as to determine the optimal number of used PCs based on cumulative variance.

#### Redundancy analysis: disentangling patterns of isolation by distance, by environment, and by barrier

We implemented a set of redundancy analyses (RDA) to disentangle the relative contribution of alternative hypotheses for explaining genetic differentiation in *A. cephalotes:* IBD, IBE, and IBB. RDA is a canonical extension of PCA, in which the principal components are constrained to be linear combinations of a set of predictors. The goal of RDA here was to identify the best ordination model that describes genetic differentiation (James et al. 2011) to better understand how spatial heterogeneity across the Andes in the Colombian Pacific and Andean regions affects patterns of gene flow in *A. cephalotes.*

For microsatellite data, a conventional RDA analysis was performed using population allele frequencies as a dependent matrix since Euclidean distance estimates involved in the analysis are directly related to *F*_ST_ values (as long as frequencies are not scaled). *mtCOI* data were analyzed through a distance-based RDA using a *Φ*_ST_-based genetic distance matrix directly. In both cases, space (geographic distances matrix), environment (climate variables), and topography (barriers) were accounted as explanatory variables. Explanatory variables were grouped into three classes according to their resulting patterns of isolation: 1) IBD, with variables representing the geographic distance between populations (space); 2) IBE, with variables determining environmental differences between populations; and 3) IBB, based on the Western range as a geographic barrier splitting populations in an allopatric manner. Because space was the only explanatory variable initially expressed as a distance matrix, it was transformed into a vector format (Oksanen et al. 2019) by PCA using the *‘pcnm’* function in the package VEGAN. Only the best explanatory PCNM components were retained. The significance of predictors was assessed using multivariate *F*-statistics with 10000 permutations using the *‘anova.cca’* function included in the package VEGAN. All explanatory variables were scaled using the *‘scale’* function in the package VEGAN. Used allele frequencies were not scaled to keep their inter-population relation with the *Φ*_ST_.

Spatial explanatory variables were tested through an IBD analysis based on RDA, followed by sequential elimination of environmental variables; only variables with | r | < 0.80 were retained (Dormann et al. 2013; Supplementary figure SF2). The remaining variables were tested through a model selection approach, comparing all possible combinations or marginal tests to be finally included in the IBE analysis. Each marginal test was compared to the null model (intercept; AIC: 1.41). These combinations of environmental variables were used to identify the best model based on the Akaike information criteria (AIC). We selected the best model for significant predictors using the *“ordistep”* function using the package VEGAN to prevent overfitting. IBB was tested using the classification of sampled populations represented by a dummy variable. Once all three models of genetic isolation were defined and tested, both IBE and IBB were tested while controlling for spatial autocorrelation through a partial RDA (conditional test). Finally, the relative contribution of IBD, IBE, and IBB to explain patterns of gene flow in *A. cephalotes* was evaluated from the final complete model, including all mechanisms of genetic isolation using a variation partitioning analysis with the function *‘varpart’* implemented in the package VEGAN. This analysis allowed us to disentangle total genetic variance according to underlying mechanisms of isolation imposed by space, environment, and barriers, as well as their relative contributions when considered together.

## RESULTS

### Genetic diversity

All microsatellite loci followed expectations from HWE or LD tests. Most loci were highly variable, with number of alleles ranging from 2 to 20 per locus (Table 2, Supplementary Table ST2). The Pacific region showed the highest allelic richness, largely due to 17 private alleles, while the Andean 1 region had the lowest allelic richness with four private alleles. Most genetic diversity parameters were significantly higher for the Pacific region, except for He estimates (Figure 2A, Supplementary Table ST4). An excess of heterozygotes was teceted in the PPY population (IAM model: T= 3.40, *P* < 0.01; SSM model: T= 1.92, *P* = 0.03) while allele frequencies were shifted from the L-shaped distribution. A putative bottleneck was supported by the low number of alleles per locus detected in this population (range: 1 to 6). Inbreeding coefficients were not significantly different from zero, suggesting random mating in all populations (Table 2).

**Figure 2.**
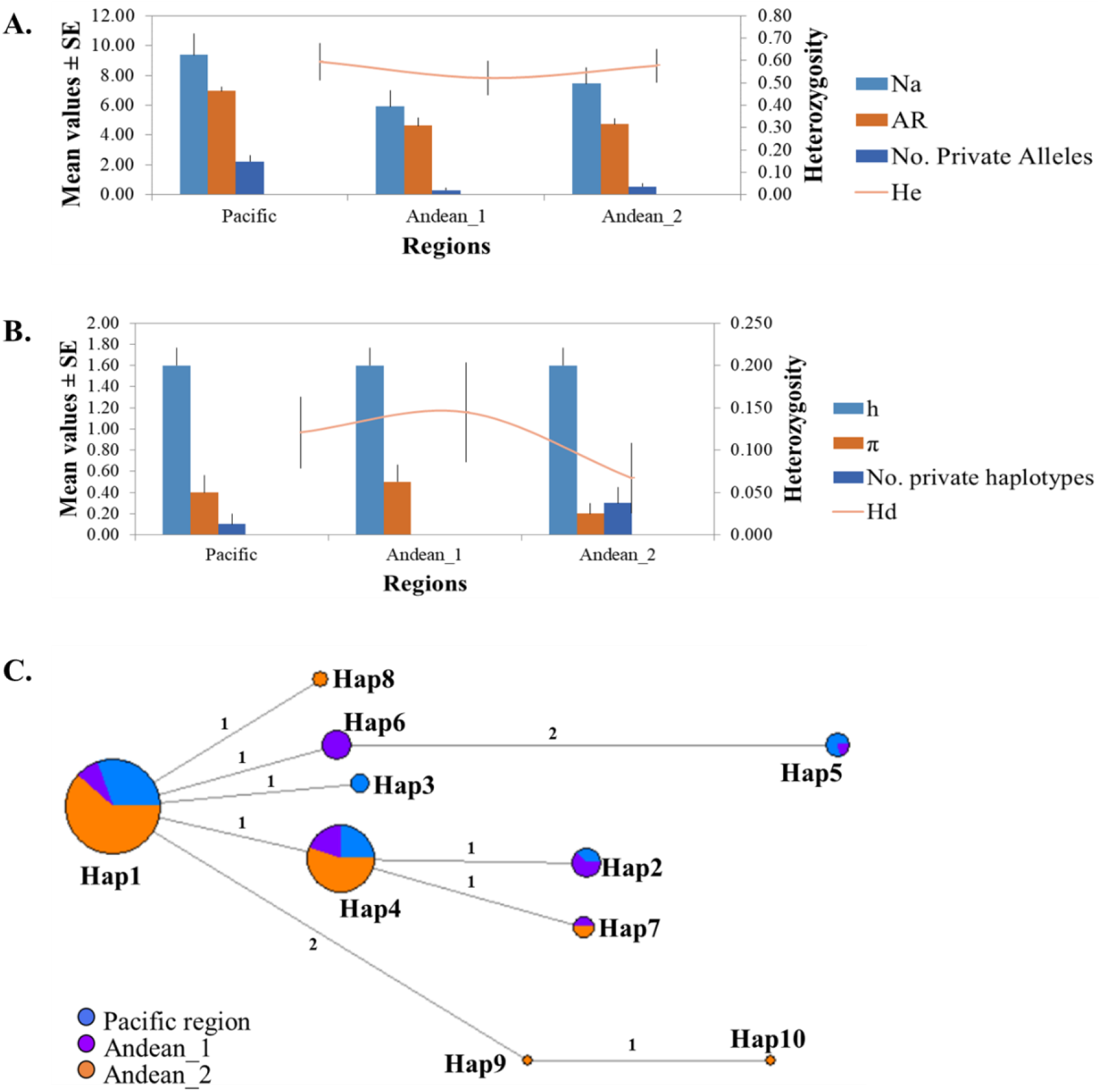
Patterns of genetic diversity and haplotype network for *A. cephalotes* sampled in three different regions from Colombia. **A.** Genetic diversity parameters for microsatellite markers (N_a_. mean number of different alleles; A_R_. mean allelic richness; H_E_: mean expected heterozygosity). **B.** Genetic diversity parameters for mitochondrial *mtCOI* (h: number of haplotypes; π: nucleotide diversity; H_d_. haplotype diversity). **C.** Haplotype network based on 146 sequences of the *A. cephalotes* mitochondrial *COI* gene. Numbers connecting lines indicate the number of mutations.

A 307 bp *mtCOI* gene fragment was successfully sequenced in 146 individuals. Ten nucleotide sites (3.3%) were polymorphic and defined 10 different haplotypes that differ in 1-3 positions (0.4% sequence divergence). The two most common haplotypes Hap1 (frequency= 0.51) and Hap4 (frequency = 0.27) were found in almost all sampling locations. One haplotype was specific to the Pacific region (Hap3), one to the Andean 1 (Hap6), and three to the Andean 2 regions (Hap8, Hap9, Hap10; Figure 2C, Supplementary Table ST5). Overall, haplotype and nucleotide diversities were Hd= 66% (range: 0.0 – 76%) and π = 0.3% (range: 0.0 – 0.4%). In contrast to the microsatellite data, Andean 1 region showed the highest diversity (Table 2, Figure 2B). However, the differences in mitochondrial diversity across regions were not significant (Figure 2B, Supplementary Table ST4).

### Population structure for nuclear and mitochondrial data

Two regional models were used in the hierarchical AMOVA following IBB and IBE scenarios (Table 3). For microsatellite data, results were similar under IBB and IBE scenarios, showing substantial differentiation among regions and populations within regions. Moreover, genetic differentiation was higher among regions when compared with that among populations and slightly higher under the IBE model when compared to the IBB model (Table 3).

**Table 3.** Hierarchical AMOVA for microsatellites and mitochondrial *mtCOI* markers. The analyses included two intermediate levels of variation following 1) Isolation by Barrier (IBB) model with populations separated by Western range (Pacific, Andean), and 2) Isolation by Environment (IBE) with populations classified into three regions following environmental conditions (Pacific, Andean 1, Andean 2). Proportion of variation (%) and associated *F*_ST_ and *Φ* indicates given for each hierarchical level.

| Model | Source of variation | Nuclear DNA |  |  | Mitochondrial DNA |  |  |
| --- | --- | --- | --- | --- | --- | --- | --- |
| | | $F_{ST}$ | | | $\Phi_{ST}$ | | |
| | | df | % | $F_{ST}$ | df | % | $\Phi$ |
| 1) IBB: Pacific populations and Andean populations | Regions | 1 | 2 | $F_{CT}: 0.02^*$ | 1 | 0 | $\Phi_{CT}: -0.08$ |
| | Pops (regions) | 8 | 9 | $F_{SC}: 0.09^*$ | 8 | 33 | $\Phi_{SC}: 0.33^*$ |
| | Within pops | 296 | 90 | $F_{ST}: 0.10^*$ | 136 | 67 | $\Phi_{ST}: 0.28^*$ |
|  | Total | 305 | 100 |  | 145 | 100 |  |
| 2) IBE: Pacific, Andean 1 and Andean 2 populations | Regions | 2 | 4 | $F_{CT}: 0.04^*$ | 2 | 0 | $\Phi_{CT}: -0.06$ |
| | Pops (regions) | 7 | 7 | $F_{SC}: 0.07^*$ | 7 | 33 | $\Phi_{SC}: 0.34^*$ |
| | Within pops | 296 | 89 | $F_{ST}: 0.11^*$ | 136 | 67 | $\Phi_{ST}: 0.30^*$ |
|  | Total | 305 | 100 |  | 145 | 100 |  |

Pairwise *F*_ST_ estimates between populations ranged from 0.01 to 0.28 (mean = 0.10) and were significant after Bonferroni corrections except for the BVT-QBD comparison in the Pacific region (Figure 3A). Pairwise comparisons between regions under the IBE model were significant, showing the highest differentiation values for comparisons involving the Andean 1 region (Pacific-Andean1: 0.04; Pacific-Andean2: 0.02; Andean1-Andean2: 0.10), consistent with the DAPC analysis (Figure 4A). However, regional comparisons under the IBB model were also significant (*F*_ST_ = 0.04).

**Figure 3.**
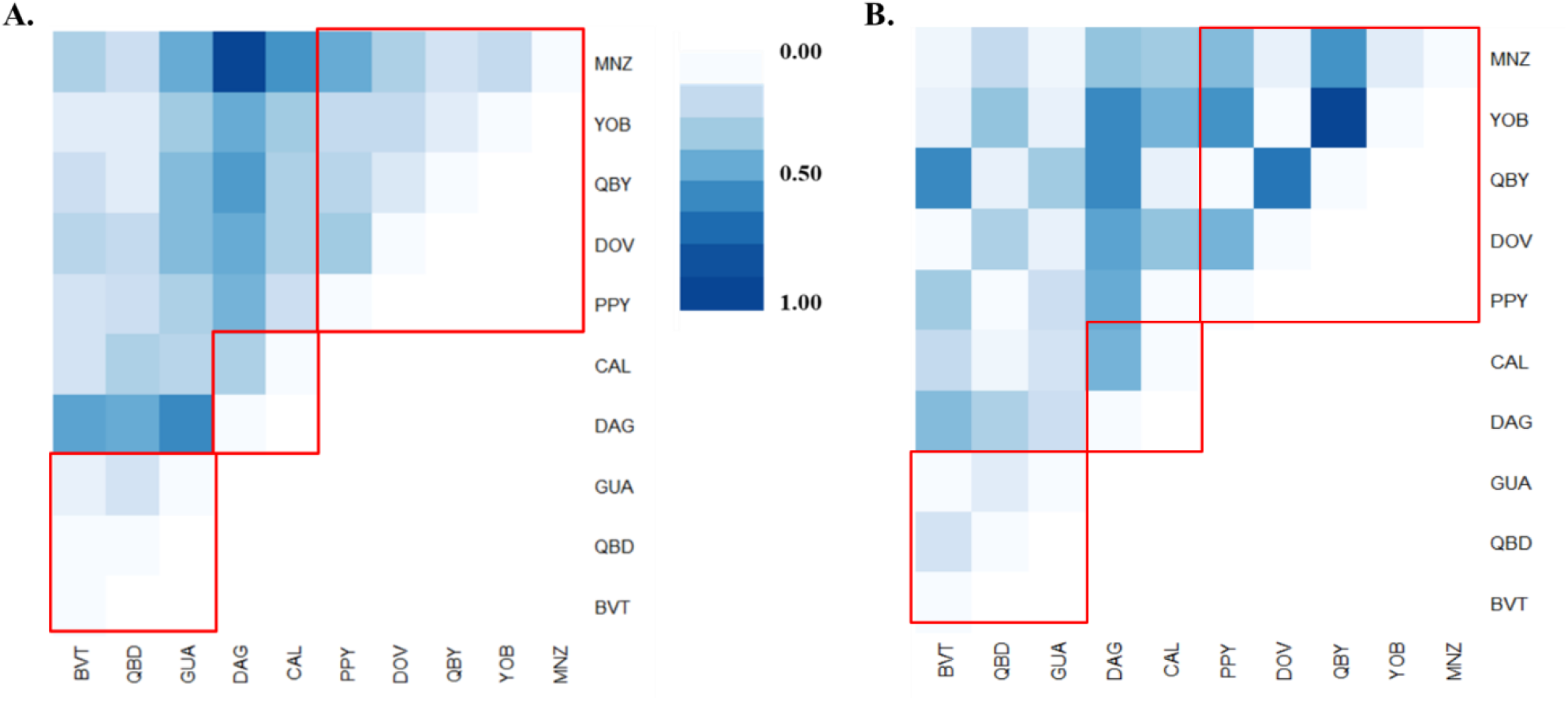
Heatmap for pairwise *F*_ST_ comparisons of *Atta cephalotes* populations. **A.** *F*_ST_ comparisons based on nuclear microsatellites and, **B.** *Φ*_ST_ comparisons based on *mtCOI.* Red squares show comparisons intra-regions.

**Figure 4.**
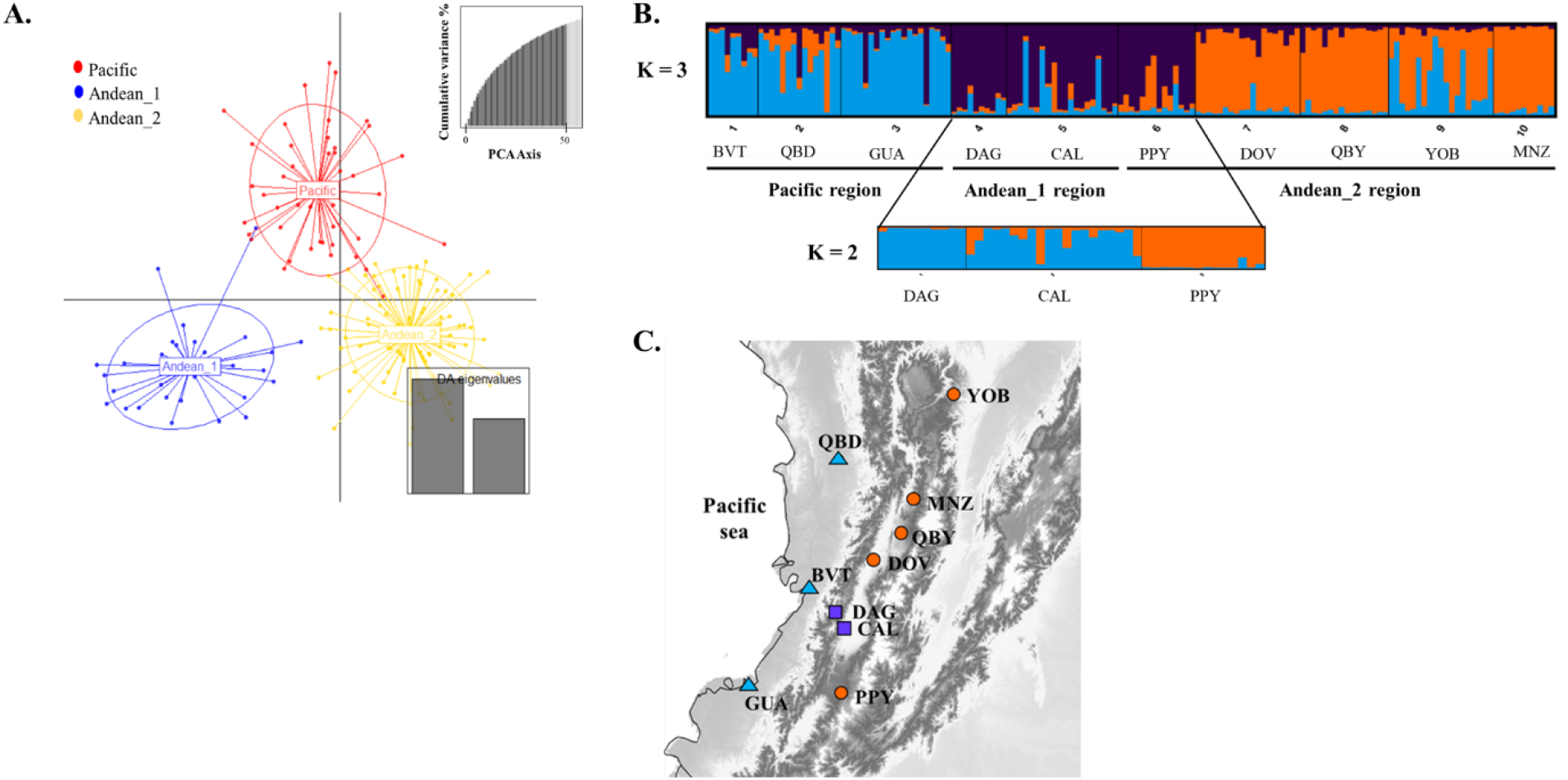
Population structure of *Atta cephalotes* as was obtained from STRUCUTRE and DAPC analyses. **A.** Multilocus genotype clustering of *A. cephalotes* populations, using STRUCTURE and following Evanno et al. (2005). Each region is characterized by a color and each location is represented by a vertical bar. Locations are organized by region starting with Pacific (BVT – GUA) followed by Andean_1 (DAG and CAL), and Andean_2 (PPY – MNZ). A different simulation was run for the south cluster. **B.** Map for sampling localities according to STRUCTURE clustering: Pacific (BVT, QBD and GUA), Andean south (DAG, CAL and PPY) and Andean north (DOV, QBY, YOB and MNZ). **C.** Discriminant analysis of principal components (DAPC) scatterplot between samples. Key describes the colors attributed to each region and inertia ellipses describe the general distribution of points. Eigenvalues for each PC axis are shown (PC1, vertical; PC2, horizontal). The number of PCA axes retained in each DAPC analyses is shown in the bottom-right inset (gray bars).

Hierarchical AMOVA for *mtCOI* data inidcated that populations within regions harbored 34% of the genetic variance, but no region effect was detected neither under IBB nor IBE models (Table 3). Pairwise *Φ*_ST_ values estimated among populations ranged from 0.00 to 0.86 and, 56% of these comparisons were significant (Figure 3B). Moreover, DAPC clustering analysis also failed to group populations by regions (Supplementary figure SF3), supporting AMOVA results.

### Clustering analyses

Evidence for hierarchical population structure in microsatellites data was also found from STRUCTURE runs. The Δ*K ad hoc* statistic by Evanno et al. (2005) indicated an optimal *K* = 3 [Pr(X / *K*: 2) = −5061.73] as the uppermost hierarchical level (Supplementary figure SF1). These three genetic clusters correspond to the individuals belonging to the three regions originally considered under the IBE model, except for the PPY population from the Andean 2 region. The first group (blue in Figure 4B) clustered most samples from the Pacific region. The second group (purple in Figure 4B) clustered most samples from Andean 1 region, as well as most samples from PPY Andean 2 region (PPY). The third group (orange in Figure 4B) clustered most samples from Andean 2 region except for PPY. This pattern suggests a hierarchical north-south structure in the Andean region rather than a climate-related structure tested through the IBE-based AMOVA (Figure 4C) and suggests IBD influencing regional differentiation for the Andean regions rather than, or in addition to IBE.

To investigate further the role of IBD in the genetic structure of *A. cephalotes*, we performed a new AMOVA by moving PPY from the Andean 2 to Andean 1 region. However, the results were similar to those under the original IBE scenario (results not shown). We also run another STRUCTURE analysis, including only samples from Andean 1 region and PPY population (southern Andean cluster). This analysis revealed two genetic groups (K = 2), where PPY formed a sole cluster (Figure 4B).

Results for the IBE model were supported by the DAPC analysis, which clustered defined regions based on environmental variables. The first two principal components of the DAPC analysis explained 77.1% of the variance in allele frequencies (50 PCs retained; Figure 4A). In this case, the first principal component clustered Pacific and Andean regions, while the second principal component reflects the differentiation between Andean 1 and Andean 2 regions (Figure 4A).

### RDA analysis – Alternative scenarios of genetic differentiation: IBD, IBE, and IBB

Significant IBD was detected from RDA analysis for microsatellite data, where the geographic distance expressed as the first axis of the PCNM analysis (PCNM1) contributed to the genetic structure of *A. cephalotes* based on model selection. This suggests that distance is an important determinant of genetic structure in *A. cephalotes* (Figure 5). These results agree with those obtained from STRUCTURE software, suggesting that genetic differentiation in the Andean regions can also be explained by an IBD pattern (north to south) and not exclusively by IBE across regions (Figure 4B). IBD results followed different patterns between nuclear and mitochondrial markers since IBD was not detected from *mtCOI* data.

**Figure 5.**
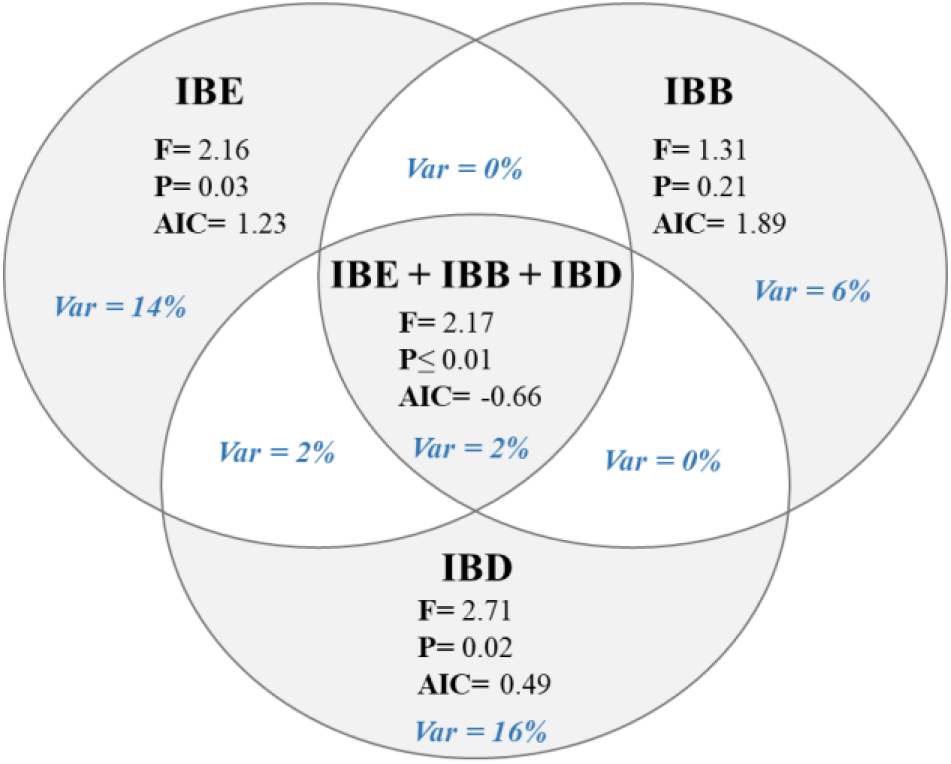
Variation partitioning and model comparison of isolating (RDA) mechanisms driving genetic nuclear differentiation (*F*_ST_) in *Atta cephalotes*. Variation partitioning analysis based on RDA results of the full model (IBE + IBB + IBD) into environmental (IBE: temperature and precipitation), barrier (IBB: Andes) and spatial (IBD: geographic distance) components. Each circle represents the variation explained for each mechanism, and their overlap the fraction of shared variation. Model selection and its significance are indicated by AIC and *P*-values. The IBE model remained significant after controlling for IBD (Conditioned model: F: 1.66, *P*: 0.04, AIC: 0.09), while IBB was not significant when accounting for IBD (Conditioned model: F: 1.49, *P*: 0.11, AIC: 0.56).

Although alternative models used for regional differentiation suggested a stronger influence of IBE, based on its higher regional genetic differentiation when compared to IBB or IBD, all models were significant (Table 3). This was also observed from RDA results, where two environmental variables (temperature and precipitation) and the barrier variable (Andes classification) were significantly associated with genetic divergence (Table 4, Figure 5). Moreover, a significant contribution of alternative models of genetic differentiation was only observed from microsatellite data (Supplementary Table ST6). The optimal model including all mechanisms of genetic isolation as tested for IBD, IBE, and IBB, accumulated up to 33% of explained variation, which remained marginally significant after accounting for IBD in conditional tests (Figure 5).

**Table 4.** Results of model selection as tested through the RDA analysis under IBD, IBE and IBB scenarios of genetic divergence.

| MODEL | F | P | AIC |
| --- | --- | --- | --- |
| **One model** |  |  |  |
| IBD model | 1.17 | 0.30 | 1.33 |
| IBD model selected (PCNM1) | 2.71 | 0.01* | 0.49 |
| IBE model (T+ H+ P+ E+ C) | 1.46 | 0.07 | 1.65 |
| IBE model selected (T+ P) | 1.81 | 0.03* | 1.23 |
| IBB model (Andes) | 1.31 | 0.22 | 1.89 |
| **Two combined models** |  |  |  |
| IBE-IBB (T+ P+ Andes) | 1.69 | 0.03* | 1.28 |
| IBE-IBD (T+ P+ PCNM1) | 2.16 | $\leq 0.01^*$ | 0.09 |
| IBB-IBD (Andes + PCNM1) | 2.17 | $\leq 0.01^*$ | 0.58 |
| **Three combined models** |  |  |  |
| Whole model (IBD+IBE+IBB) | 2.17 | $\leq 0.01^*$ | -0.66 |
| **Conditioned models by PCNM1** |  |  |  |
| IBE model (T+ P) ~ PCNM1 | 1.65 | 0.04* | 0.09 |
| IBB model (Andes) ~ PCNM1 | 1.48 | 0.13 | 0.58 |
| Whole model (IBD+IBE+IBB) ~ PCNM1 | 1.74 | 0.02* | -0.66 |

T: Temperature; H: Humidity; P: Precipitation; E: Elevation; C: Climate. (\*) Indicates significant values, $P \leq 0.01$ . AIC null model: 1.41.

Variation partitioning for the full model, including all alternative models of genetic isolation, showed that IBD and IBE explained the highest proportion of genetic variation relative to IBB (Figure 5). Interaction effects between major mechanisms of genetic isolation were low for the triple interaction (IBD + IBE + IBB = 2%) and were not significant for paired interactions (Figure 5). The higher fraction of unexplained variation (residuals) might reflect the occurrence of genetic drift within populations, which is not associated with the explanatory variables.

## DISCUSSION

The complex landscape across the Andean uplift is an important barrier isolating populations and increasing genetic divergence on both sides of the mountains (Antonelli et al. 2009; Hoorn et al. 2010; Salgado-Roa et al. 2018). The Western range of the Andes in Colombia is one of the most biodiverse ecosystems in the Neotropics (Kattan et al. 2004; Salgado-Roa et al. 2018) and the most complex landscape within the distribution of the leaf-cutting ant *A. cephalotes*. By integrating hierarchical population structure with models of isolation by distance (IBD), by environment (IBE), and by barrier (IBB), we explored the role of the Western range on the distribution of genetic variation of *A. cephalotes.* Here, we demonstrated that the environmental heterogeneity imposed by the Andean uplift has highly influenced population structure in *A. cephalotes*.

### Genetic diversity

Our results indicate that populations of *A. cephalotes* are highly variable at nuclear markers, with significant genetic differentiation at region and population levels. In contrast, populations are highly differentiated for mitochondrial *COI* gene, but no region effect was detected with this marker. Ten *mtCOI* haplotypes were detected in 146 nests analyzed, showing low nucleotide and moderate haplotype diversities, with most populations dominated by the same haplotype (Hap1). As expected from a common biogeographical history, similar *mtCOI* patterns have been detected in the leaf-cutting ant *A. colombica* (Helmkampf et al. 2008), where six *mtCOI* haplotypes and relatively low levels of nucleotide diversity (0.1% population-wide) were detected across 20 colonies, with most specimens sharing the same haplotype. However, these results were obtained from a small sample size in a small area, and despite that several other genetic studies have been reported in Attini (*e.g., Acromyrmex*) (Diehl et al. 2001, 2002; Cantagalli et al. 2013; Pinheiro dos Reis et al. 2014), the nature of markers and sampling prevents us from meaningful comparisons on diversity estimates.

### Spatial differentiation at regional and local levels

We detected significant genetic differentiation across regions from nuclear data, as evidenced from AMOVA analyses and clustering methods, suggesting a potential role of the Andean landscape in restricting *A. cephalotes*’ gene flow. We tested alternative scenarios causing regional differentiation considering IBE as model of divergence, as well as a geographic barrier imposed by the Western range under an IBB model.

Regional differentiation from microsatellite data was higher under the IBE than the IBB model, suggesting a more prominent role of climate restricting gene flow as opposed to a geographic barrier in *A. cephalotes.* On the other hand, STRUCTURE results support a regional differentiation with mixed effects between IBE and IBD, evidencing a spatial North-South pattern for Andean populations rather than a regional climate model. However, this IBD pattern was likely due to a single population. PPY was initially classified within the Andean 2 region but clustered mostly (Q > 0.95) with the geographically closer Andean 1 region. Nevertheless, this pattern occurs only in the Andean region, suggesting that population differentiation in this particular case results from dispersal distance rather than environmental conditions or a geographic barrier imposed by the Western range. Medium to long dispersal distances have been reported for *A. cephalotes*, with queens flying up to 50 km during nuptial flights (Cherrett 1968; Helms 2018). This distance is at least one order of magnitude smaller than the inter-population distance in this study, but it is also probably an underestimate as it was measured in a small island (Helms 2018). Moreover, only a small number of individuals are needed to homogenize population allele frequencies and gene flow can also be conveyed stepwise. Although genetic variance at region level suggested IBE, all models of hierarchical differentiation were significant for RDA analysis, suggesting that all three leading causes of genetic isolation are likely contributing to the observed patterns of gene flow.

Significant nuclear differentiation at the population level suggests that additional features besides proximity, environment and major dispersal barriers restrict gene flow. The markedly increased variance in the PPY population is consistent with a recent population expansion signal. Whether the bottleneck signal is a product of a founder effect (Barton and Turelli 2004; Parisod et al. 2005; Matute 2013) or a recent change in effective population size cannot be determined with current data. However, the proximity and clustering of PPY to Andean 1 populations rather than Andean 2 region in STRUCTURE results, suggests a recent founder effect or an admixture between the two Andean regions.

As opposed to nuclear data, there was no regional differentiation in *mtCOI* data, but populations were significantly differentiated within the regions in both IBE and IBB models. Muñoz-Valencia et al. (2021) used *mtCOI* data to study the genetic structure of *A. cephalotes* from a larger geographic range covering most of the distribution of the species in South and Central America. They found significant genetic differentiation both at a regional scale and among populations within major regions. Together, these results show that the Eastern range of the Andes Mountains appears to be the major dispersal barrier driving regional differentiation, while the West range forms a relatively homogenous biogeographic area.

### Disentangling patterns of IBD, and IBE, IBB

Our RDA framework provides strong evidence for restricted gene flow in *A. cephalotes* across the Western Mountain range of Colombian Andes by all mechanisms tested. Furthermore, we found that explained genetic variation was maximized when all spatial and environmental variables associated with these mechanisms were included in the model. These multifactorial patterns are often expected from complex and heterogeneous environments (Shafer and Wolf 2013; Sexton et al. 2014), such as in the Colombian Andes (Kattan et al. 2004; Pérez-Escobar et al. 2017; Salgado-Roa et al. 2018). Regions classified according to climate variation and geographic barriers follow IBE but also include IBB, supporting the higher differentiation estimate (*F*_CT_) obtained under the IBE model and the significance when both models are included in the RDA (IBE + IBB). Migration through low elevation passes in the Western Mountain range (Hernández-Camacho 1992; Kattan et al. 2004) would allow gene flow between regions, explaining why such a major geographic barrier did not have a more pronounced effect. Spatial and environmental variables presumably associated with restriction to gene flow often suffer from spatial autocorrelation, challenging the differentiation of their underlying causes (Crispo et al. 2006; Edwards et al. 2012; Wang et al. 2013). Our RDA framework accounted for this issue demonstrating strong IBE even after controlling for IBD; while IBB was not significant, even when ignoring IBD. Therefore, temperature and precipitation appear to be the leading causes of IBE in *A. cephalotes.*

The significant effect of the environment on the genetic structure of *A. cephalotes* populations indicates that they have the potential to adapt to local environmental conditions. This may occur when processes such as sex-biased dispersal (Edelaar and Bolnick 2012), or selection against migrants (Wright 1943; Hendry 2004; Weber et al. 2016) decrease the rate of gene flow as has been observed in several species (Lee and Mitchell-Olds 2011; Wang et al. 2013; De Queiroz et al. 2017). Moreover, geographic distance and barriers in between can restrict gene flow by reducing dispersal efficiency, more associated with local genetic drift (Clémencet et al. 2005; Cross et al. 2016; Noguerales et al. 2016; De Queiroz et al. 2017; Smith et al. 2018), leading to combined patterns of IBD and IBB. These complex patterns have been detected, for example, in the Amazonian common sardine fish *Triportheus albus* (De Queiroz et al. 2017). However, disentangling individual environmental drivers was not possible due to strong correlations across environmental variables (Crispo et al. 2006; Edwards et al. 2012; Wang et al. 2013). The underlying causes of genetic isolation of populations have been rarely explored in leaf-cutting ants (Branstetter et al. 2017). In *A. sexdens rubipilosa*, geographic separation of populations did not explain population divergence alone (Cantagalli et al., 2013), and significant IBD and biome fragmentation imposed physical barriers to gene flow in *A. robusta* (Pinheiro dos Reis et al. 2014). All these results together strongly suggest that investigated isolating mechanisms are not mutually exclusive (Crispo et al. 2006; Edwards et al. 2012; Wang et al. 2013), and are important in the evolution of Neotropical species.

Agricultural activities have significantly contributed to the colonization processes of leaf-cutting ants (de Carvalho Cabral 2015). Conversion of the natural habitat into cultivated land may favor population growth and dispersal of leaf-cutting ants (Montoya-Lerma et al. 2012; Schowalter and Ring 2017). Queens have been shown to prefer nesting in open areas rather than closed forests in *Atta laevigata* (Vasconcelos 1997). Our exceptional study population PPY is located in a large city (Popayán). It is feasible that the city infrastructure and constant human interference rather than natural dispersal associated with nuptial flights have influenced its genetic structure. This pattern is more compatible with human-mediated dispersal (Zheng et al. 2018), where environmental changes associated with human expansion appear to promote population growth in leaf-cutting ants (Montoya-Lerma et al. 2012; Siqueira et al. 2017). However, PPY is most likely an exception from a general large-scale migration process in *A. cephalotes*.

The Andean uplift appears to modulate population structure of *A. cephalotes* in a more complex manner than previously thought. We previously demonstrated that the Eastern Mountain range of the Andean uplift in Colombia plays a major role as a geographic barrier to historical gene flow, restricting *A. cephalotes*’ dispersion from north to south (Muñoz-Valencia et al. 2021). By exploring a finer scale across the Western Mountain range and incorporating neutral genetic markers with environmental variables, we clearly show that observed genetic differentiation in *A. cephalotes* is mostly affected by a combined multifactorial effect of different isolating mechanisms mediated by the landscape complexity. Together, these results suggest that studies aiming to explore historical gene flow across the Andes Mountains should be interpreted with caution as the complexity and history of the landscape can dramatically influence results at different scales.

## ACKNOWLEDGEMENTS

We thank Kirsi Kähkönen, the ESB (Evolution, Sociality, Behavior) research group, and everyone in the Molecular Ecology and Systematics laboratory (University of Helsinki) for helping to develop the DNA microsatellite loci. We also thank you, Sandra Milena Valencia Giraldo, Andrea López-Peña, and Glever Alexander Vélez-Martínez for helping with DNA extraction. This work was funding by Vice-rectoría de Investigaciones, Universidad del Valle, Cali, Colombia (grant number: CI71067); and COLCIENCIAS National Program of PhD (grant number: 617-2013).

## AUTHORS’ CONTRIBUTIONS

VMV participated in the conceptualization, ant sampling, data analysis, original draft, reviewing, and editing of the manuscript. JML participated in conceptualization, supervision, original draft, reviewing, and editing the manuscript. PS participated in the method validation, data analysis, reviewing, and editing of the manuscript. FD participated in conceptualization, data analysis, original draft, reviewing, and editing the manuscript. All authors read and approved the final version of the manuscript.

## Notes

### Competing Interest Statement

The authors have declared no competing interest.

## REFERENCES

Antonelli A, Quijada-Mascareñas A, Crawford AJ, et al (2009) Molecular studies and phylogeography of Amazonian tetrapods and their relation to geological and climatic models. In: Hoorn C, Wesselingh F (eds) Amazonia, landscape and species evolution: A look into the past. Blackwell Publishing Ltd, London, UK, pp 386–404

Barton NH, Turelli M (2004) Effects of genetic drift on variance components under a general model of epistasis. Evolution (N Y) 58:2111–2132. doi: 10.1111/j.0014-3820.2004.tb01591.x

Branstetter MG, Jesovnik A, Sosa-Calvo J, et al (2017) Dry habitats were crucibles of domestication in the evolution of agriculture in ants. Proc R Soc B 284:20170095

Cadena CD, Pedraza CA, Brumfield RT (2016) Climate, habitat associations and the potential distributions of Neotropical birds: Implications for diversification across the Andes. Rev la Acad Colomb Ciencias Exactas, Físicas y Nat 40:275. doi: 10.18257/raccefyn.280

Cantagalli LB, Mangolin CA, Ruvolo-Takasusuki MCC (2013) Population genetics of Atta sexdens rubropilosa (Hymenoptera: Formicidae). Acta Biológica Colomb 18:179–190

Carro B, Quintela M, Ruiz JM, Barreiro R (2019) Wave exposure as a driver of isolation by environment in the marine gastropod Nucella lapillus. Hydrobiologia 839:51–69. doi: 10.1007/s10750-019-03993-5

Chambers JM, Freeny A, Heiberger RM (1992) Analysis of variance; designed experiments. In: Chambers JM, Hastie TJ (eds) Statistical models. Wadsworth & Brooks/Cole

Chapuis MP, Estoup A (2007) Microsatellite null alleles and estimation of population differentiation. Mol Biol Evol 24:621–631. doi: 10.1093/molbev/msl191

Chen D, Chen HW (2013) Using the Köppen classification to quantify climate variation and change: An example for 1901-2010. Environ Dev 6:69–79. doi: 10.1016/j.envdev.2013.03.007

Cherrett J (1968) A flight record for queens of Atta cephalotes L. (Hym. Formicidae). Entomol’s Mon Mag 104:255–256

Clémencet J, Viginier B, Doums C (2005) Hierarchical analysis of population genetic structure in the monogynous ant Cataglyphis cursor using microsatellite and mitochondrial DNA markers. Mol Ecol 14:3735–3744. doi: 10.1111/j.1365-294X.2005.02706.x

Crispo E, Bentzen P, Reznick DN, et al (2006) The relative influence of natural selection and geography on gene flow in guppies. Mol Ecol 15:49–62. doi: 10.1111/j.1365-294X.2005.02764.x

Cross TB, Naugle DE, Carlson JC, Schwartz MK (2016) Hierarchical population structure in greater sage-grouse provides insight into management boundary delineation. Conserv Genet 17:1417–1433. doi: 10.1007/s10592-016-0872-z

De-Silva DL, Mota LL, Chazot N, et al (2017) North Andean origin and diversification of the largest ithomiine butterfly genus. Sci Rep 7:1–17. doi: 10.1038/srep45966

de Carvalho Cabral D (2015) Into the bowels of tropical earth: Leaf-cutting ants and the colonial making of agrarian Brazil. J Hist Geogr 50:92–105. doi: 10.1016/j.jhg.2015.06.014

De Queiroz LJ, Torrente-Vilara G, Quilodran C, et al (2017) Multifactorial genetic divergence processes drive the onset of speciation in an Amazonian fish. PLoS One 12:1–27. doi: 10.1371/journal.pone.0189349

Della Lucia TM, Gandra LC, Guedes RN (2014) Managing leaf-cutting ants: Peculiarities, trends and challenges. Pest Manag Sci 70:14–23. doi: 10.1002/ps.3660

Diehl E, Cavalli-molina S, Mellender de Araujo A (2002) Isoenzyme variation in the leaf-cutting ants Acromyrmex heyeri and Acromyrmex striatus (Hymenoptera, Formicidae). Genet Mol Biol 25:173–178

Diehl E, de Araújo AM, Cavalli-Molina S (2001) Genetic variability and social structure of Colonies in Acromyrmex heyeri and A. striatus (Hymenoptera: Formicidae). Braz J Biol 61:667–678. doi: 10.1590/S1519-69842001000400017

Dormann CF, Elith J, Bacher S, et al (2013) Collinearity: A review of methods to deal with it and a simulation study evaluating their performance. Ecography (Cop) 36:027–046. doi: 10.1111/j.1600-0587.2012.07348.x

Earl DA, vonHoldt BM (2012) STRUCTURE HARVESTER: a website and program for visualizing STRUCTURE output and implementing the Evanno method. Conserv Genet Resour 4:359–361. doi: 10.1007/s12686-011-9548-7.

Edelaar P, Bolnick DI (2012) Non-random gene flow: An underappreciated force in evolution and ecology. Trends Ecol Evol 27:659–665. doi: 10.1016/j.tree.2012.07.009

Edwards DL, Keogh JS, Knowles LL (2012) Effects of vicariant barriers, habitat stability, population isolation and environmental features on species divergence in the south-western Australian coastal reptile community. Mol Ecol 21:3809–3822. doi: 10.1111/j.1365-294X.2012.05637.x

Evanno G, Regnaut S, Goudet J (2005) Detecting the number of clusters of individuals using the software STRUCTURE: A simulation study. Mol Ecol 14:2611–2620. doi: 10.1111/j.1365-294X.2005.02553.x

Excoffier L, Lischer HEL (2010) Arlequin suite ver 3.5: A new series of programs to perform population genetics analyses under Linux and Windows. Mol Ecol Resour 10:564–567. doi: 10.1111/j.1755-0998.2010.02847.x

Excoffier L, Smouse PE, Quattro JM (1992) Analysis of molecular variance inferred from metric distances among DNA haplotypes: Application to human mitochondiral DNA restriction data. Genetics 131:479–491. doi: 10.1007/s00424-009-0730-7

Fernández F, Castro-Huertas V, Serna F (2015) Hormigas cortadoras de hojas de Colombia: Acromyrmex & Atta (Hymenoptera: Formicidae), Monografía. Instituto de Ciencias Naturales, Universidad Nacional de Colombia, Bogotá

Goudet J (1995) FSTAT (Version 1.2): A Computer program to Calculate F-Statistics. J. Hered. 485–486

Goudet J (2001) FSTAT, a program to estimate and test gene diversities and fixation indices (version 2.9.3)

Haffer J (2008) Hypotheses to explain the origin of species in Amazonia. Brazilian J Biol 68:917–947. doi: 10.1590/S1519-69842008000500003

Hakala SM, Seppä P, Helanterä H (2019) Evolution of dispersal in ants (Hymeno ptera: Formicidae): a review on the dispersal strategies of sessile superorganisms Sanja. Myrmecological News 29:35–55. doi: 10.25849/myrmecol.news

Helmkampf M, Gadau J, Feldhaar H (2008) Population- and sociogenetic structure of the leaf-cutter ant Atta colombica (Formicidae, Myrmicinae). Insectes Soc 55:434–442. doi: 10.1007/s00040-008-1024-3

Helms J (2018) The flight ecology of ants (Hymenoptera: Formicidae). Myrmecological News 26:19–30. doi: 10.25849/myrmecol.news

Hendry AP (2004) Selection against migrants contributes to the rapid evolution of ecologically dependent reproductive isolation. Evol Ecol Res 6:1219–1236

Hermelin M (2015) Landscapes and landforms of Colombia. Springer International Publishing

Hernández-Camacho J (1992) Caracterización geográfica de Colombia. In: Halffter G (ed) La diversidad biológica de Iberoamérica I, Programa I. Acta Zoológica Mexicana, pp 45–52

Hölldobler B, Wilson EO (2011) The leafcutter ants: civilization by instinct, 1st ed. W. W. Norton & Company, Nueva York, NY

Hoorn C, Wesselingh FP, Ter Steege H, et al (2010) Amazonia through time: Andean uplift, climate change, landscape evolution, and biodiversity. Science (80-) 330:927–931. doi: 10.1126/science.1194585

James PMA, Coltman DW, Murray BW, et al (2011) Spatial genetic structure of a symbiotic beetle-fungal system: Toward multi-taxa integrated landscape genetics. PLoS One 6:1–11. doi: 10.1371/journal.pone.0025359

Jombart T, Devillard S, Balloux F (2010) Discriminant analysis of principal components: a new method for the analysis of genetically structured populations. BMC Genet 11:1–15. doi: 10.1371/journal.pcbi.1000455

Kattan GH, Franco P, Rojas V, Morales G (2004) Biological diversification in a complex region: A spatial analysis of faunistic diversity and biogeography of the Andes of Colombia. J Biogeogr 31:1829–1839. doi: 10.1111/j.1365-2699.2004.01109.x

Kopelman NM, Mayzel J, Jakobsson M, et al (2015) CLUMPAK: a program for identifying clustering modes and packaging population structure inferences across K. Mol Ecol Resour 15:1179–1191. doi: 10.1111/1755-0998.12387.CLUMPAK

Köppen W (1884) Die Wärmezonen der Erde, nach der Dauer der heissen, gemässigten und kalten Zeit und nach der Wirkung der Wárme auf die organische Welt betrachtet (The thermal zones of the earth according to the duration of hot, moderate and cold periods and to the impac. Meteorol Zeitschrift 1:215–226. doi: 10.1127/0941-2948/2011/105

Kronauer DJC, Hölldobler B, Gadau J (2004) Phylogenetics of the new world honey ants (genus Myrmecocystus) estimated from mitochondrial DNA sequences. Mol Phylogenet Evol 32:416–421. doi: 10.1016/j.ympev.2004.03.011

Lagomarsino LP, Condamine FL, Antonelli A, et al (2016) The abiotic and biotic drivers of rapid diversification in Andean bellflowers (Campanulaceae). New Phytol 210:1430–1442. doi: 10.1111/nph.13920

Lee C-R, Mitchell-Olds T (2011) Quantifying effects of environmental and geographical factors on patterns of genetic differentiation. Mol Ecol 20:4631–4642. doi: 10.5061/dryad.6rs51

Librado P, Rozas J (2009) DnaSP v5: A software for comprehensive analysis of DNA polymorphism data. Bioinformatics 25:1451–1452. doi: 10.1093/bioinformatics/btp187

Luebert F, Weigend M (2014) Phylogenetic insights into Andean plant diversification. Front Ecol Evol 2:1–17. doi: 10.3389/fevo.2014.00027

Manel S, Holderegger R (2013) Ten years of landscape genetics. Trends Ecol Evol 28:614–621. doi: 10.1016/j.tree.2013.05.012

Manel S, Schwartz MK, Luikart G, Taberlet P (2003) Landscape genetics: combining landscape ecology and population genetics. Trends Ecol Evol 18:189–197. doi: 10.1016/S0169-5347(03)00008-9

Matute DR (2013) The role of founder effects on the evolution of reproductive isolation. J Evol Biol 26:2299–2311. doi: 10.1111/jeb.12246

Meirmans PG (2015) Seven common mistakes in population genetics and how to avoid them. Mol Ecol 24:3223–3231

Meirmans PG (2006) Using the AMOVA framework to estimate a standardized genetic differentiation measure. Evolution (N Y) 60:2399–2402

Meirmans PG, Hedrick PW (2011) Assessing population structure: FSTand related measures. Mol Ecol Resour 11:5–18. doi: 10.1111/j.1755-0998.2010.02927.x

Montoya-Lerma J, Giraldo-Echeverri C, Armbrecht I, et al (2012) Leaf-cutting ants revisited: Towards rational management and control. Int J Pest Manag 58:225–247. doi: 10.1080/09670874.2012.663946

Moser JC (1967) Mating activities of Atta texana (Hymenoptera, Fromicidae). Insectes Soc XIV:295–312

Mueller UG, Ishak HD, Bruschi SM, et al (2017) Biogeography of mutualistic fungi cultivated by leafcutter ants. Mol Ecol 1–17. doi: 10.1111/mec.14431

Muñoz-Valencia V, Kähkönen K, Montoya-Lerma J, Díaz F (2020) Characterization of a New Set of Microsatellite Markers Suggests Polygyny and Polyandry in Atta cephalotes (Hymenoptera: Formicidae). J Econ Entomol 113:3021–3027. doi: 10.1093/jee/toaa200

Muñoz-Valencia V, Vélez-Matínez GA, Montoya-Lerma J, Díaz F (2021) Role of the Andean uplift as an asymmetrical barrier to gene flow in the neotropical leaf-cutting ant Atta cephalotes. Biotropica 1–14. doi: 10.1111/btp.13050

Noguerales V, Cordero PJ, Ortego J (2016) Hierarchical genetic structure shaped by topography in a narrow-endemic montane grasshopper. BMC Evol Biol 16:1–15. doi: 10.1186/s12862-016-0663-7

Oksanen J, Blanchet FG, Friendly M, et al (2019) Vegan: Community Ecology Package. R Package Version 2.2-0

Pamilo PK, Seppä P V., Helanterä HO (2016) Population genetics of wood ants. In: Stockan JA, Robinson EJH (eds) Wood ant ecology and conservation. Cambridge University Press, pp 51–80

Parisod C, Trippi C, Galland N (2005) Genetic variability and founder effect in the pitcher plant Sarracenia purpurea (Sarraceniaceae) in populations introduced into Switzerland: from inbreeding to invasion. Ann Bot 95:277–286. doi: 10.1093/aob/mci023

Peakall R, Smouse P (2012) GenAlEx 6.5: Genetic analysis in Excel. Population genetic software for teaching and research. Bioinformatics 1:6–8

Peel MC, Finlayson BL, McMahon TA (2007) Update world map of the Köppen-Geiger climate classification. Hydrol Earth Syst Sci 11:1633–1644. doi: 10.1002/ppp.421

Pérez-Escobar OA, Gottschling M, Chomicki G, et al (2017) Andean mountain building did not preclude dispersal of lowland epiphytic orchids in the Neotropics. Sci Rep 7:1–10. doi: 10.1038/s41598-017-04261-z

Pinheiro dos Reis E, Fernandes Salomão TM, de Oliveira Campos LA, Garcia Tavares M (2014) Genetic diversity and structure of Atta robusta (Hymenoptera, Formicidae, Attini), an endangered species endemic to the Restinga ecoregion. Genet Mol Biol 37:581–586. doi: 10.1590/S1415-47572014000400015

Piry S, Luikart G, Cornuet JM (1999) BOTTLENECK: A computer porgram for detecting recen reductions in the effective population size using allele frequency data. J Hered 90:502–503. doi: 10.1093/jhered/90.4.502

Pritchard JK, Stephens M, Donnelly P (2000) Inference of population structure using multilocus genotype data. Genetics 155:945–959. doi: 10.1111/j.1471-8286.2007.01758.x

Ramos-Onsins SE, Rozas J (2002) Statistical properties of new neutrality tests against population growth. Mol Biol Evol 19:2092–2100. doi:Doi 10.1093/Molbev/Msl052

Rousset F (2008) GENEPOP’007: A complete re-implementation of the GENEPOP software for Windows and Linux. Mol Ecol Resour 8:103–106. doi: 10.1111/j.1471-8286.2007.01931.x

Rull V (2011) Neotropical biodiversity: timing and potential drivers. Trends Ecol Evol 26:508–513. doi: 10.1016/j.tree.2011.05.011

Salgado-Roa FC, Pardo-Diaz C, Lasso E, et al (2018) Gene flow and Andean uplift shape the diversification of Gasteracantha cancriformis (Araneae: Araneidae) in Northern South America. Ecol Evol 8:7131–7142. doi: 10.1002/ece3.4237

Schowalter TD, Ring DR (2017) Biology and management of the Texas leafcutting ant (Hymenoptera: Formicidae). J Integr Pest Manag 8:1–8. doi: 10.1093/jipm/pmx013

Seppä P (2008) Do ants (Hymenoptera: Formicidae) need conservation and does ant conservation need genetics? Myrmecological News 11:161–172

Seppä P, Gyllenstrand N, Corander J, Pamilo P (2004) Coexistence of the social types: Genetic population structure in the ant Formica exsecta. Evolution (N Y) 58:2462–2471. doi: 10.1111/j.0014-3820.2004.tb00875.x

Sexton JP, Hangartner SB, Hoffmann AA (2014) Genetic isolation by environment or distance: which pattern of gene flow is most common? Evolution (N Y) 68:1–15. doi: 10.1111/evo.12258

Shafer ABA, Wolf JBW (2013) Widespread evidence for incipient ecological speciation: a meta-analysis of isolation-by-ecology. Ecol Lett 16:940–950. doi: 10.1111/ele.12120

Simon C, Frati F, Beckenbach A, et al (1994) Evolution, weighting, and phylogenetic utility of mitochondrial gene sequences and a compilation of conserved polymerase chain reaction primers. Ann Entomol Soc Am 87:651–701. doi: 10.1093/aesa/87.6.651

Siqueira FFS, Ribeiro-Neto JD, Tabarelli M, et al (2017) Leaf-cutting ant populations profit from human disturbances in tropical dry forest in Brazil. J Trop Ecol 33:337–344. doi: 10.1017/S0266467417000311

Slatkin M (1993) Isolation by distance in equilibrium and non-equilibrium populations. Evolution (N Y) 47:264–279

Smith CC, Weber JN, Mikheyev AS, et al (2018) Landscape genomics of an obligate mutualism: discordant population structures between a leafcutter-ant and its fungal cultivars. bioRxiv 1–35. doi: 10.1101/458950

Sobel JM (2014) Ecogeographic isolation and speciation in the genus Mimulus. Am Nat 184:565–579. doi: 10.1086/678235

Sundström L, Seppä P, Pamilo P (2005) Genetic population structure and dispersal patterns in Formica ants – a review. Ann Zool Fennici 42:163–177. doi: 10.1007/BF00172934

Turchetto-Zolet A, Pinheiro F, Salgueiro F, Palma-Silva C (2013) Phylogeographical patterns shed light on evolutionary process in South America. Mol Ecol 22:1193–1213. doi: 10.1111/mec.12164

VanOosterhout C, Hutchinson WF, Wills DPM, Shipley P (2004) MICRO-CHECKER: Software for identifying and correcting genotyping errors in microsatellite data. Mol Ecol Notes 4:535–538. doi: 10.1111/j.1471-8286.2004.00684.x

Vasconcelos HL (1997) Foraging activity of an Amazonian leaf-cutting ant: Responses to changes in the availability of woody plants and to previous plant damage. Oecologia 112:370–378. doi: 10.1007/s004420050322

Wang IJ, Bradburd GS (2014) Isolation by environment. Mol Ecol 23:5649–5662. DOI: 10.1111/mec.12938

Wang IJ, Glor RE, Losos JB (2013) Quantifying the roles of ecology and geography in spatial genetic divergence. Ecol Lett 16:175–182. doi: 10.1111/ele.12025

Wasko AP, Martins C, Oliveira C, Foresti F (2003) Non-destructive genetic sampling in fish. An improved method for DNA extraction from fish fins and scales. Hereditas 138:161–165

Weber JN, Bradburd GS, Stuart YE, et al (2016) Partitioning the effects of isolation by distance, environment, and physical barriers on genomic divergence between parapatric threespine stickleback. Evolution (N Y) 71–2:342–356. doi: 10.1111/evo.13545

Wright S (1943) Isolation by distance. Genetics 28:114–138

Zheng C, Yang F, Zeng L, et al (2018) Genetic diversity and colony structure of Tapinoma melanocephalum on the islands and mainland of South China. Ecol Evol 8:5427–544. DOI: 10.1002/ece3.4065

