## Supplementary Material for "Landscape genetics across the Andes Mountains: Environmental variation drives genetic divergence in the leaf-cutting ant *Atta cephalotes*"

**Supplementary table ST1. Sampled populations of *A. cephalotes*.** Locations with their regional classifications (Pacific, Andean 1 and Andean 2 regions from Colombia) and landscape features are indicated for each population.

| Code | Location | Region | Coordinates | | Environmental variables | | | | Barrier variables | |
|---|---|---|---|---|---|---|---|---|---|---|
| | | | North | West | T (°C) | RH (%) | Prec. (mm) | Climate | Elev. (m.a.s.l.) | Andes |
| BVT | Buenaventura | Pacific | 3.88564 | -77.02562 | 26.1 | 89 | 7650 | 1 | 7 | 0 |
| QBD | Quibdó | Pacific | 5.6733056 | -76.6193889 | 26.6 | 87 | 7328 | 1 | 43 | 0 |
| GUA | Guapi | Pacific | 2.542533 | -77.866.933 | 26.1 | 87 | 4364 | 1 | 4 | 0 |
| DAG | Dagua | Andean_1 | 3.5350278 | -76.6462778 | 23.5 | 78 | 377.8 | 2 | 885 | 1 |
| CAL | Cali | Andean_1 | 3.3465278 | -76.5291111 | 25.1 | 71.1 | 1240.8 | 2 | 956 | 1 |
| PPY | Popayán | Andean_2 | 2.429983 | -76.604591 | 17.8 | 75 | 2040 | 3 | 1731 | 1 |
| DOV | Dovio | Andean_2 | 4.29509 | -76.13037 | 20.6 | 89 | 1321 | 4 | 1440 | 1 |
| QBY | Quimbaya | Andean_2 | 4.6368111 | -75.7740944 | 20.9 | 78 | 2024 | 4 | 1312 | 1 |
| YOB | Yolombó | Andean_2 | 6.567865 | -75.013725 | 21 | 68 | 2837 | 4 | 1444 | 1 |
| MNZ | Manizales | Andean_2 | 5.104877 | -75.572693 | 18 | 80 | 1878 | 3 | 2117 | 1 |

T (°C): average of annual temperature; RH (%): average of annual humidity; Prec. (mm): average of annual precipitation; Elev. (m.a.s.l.): elevation relative to sea level. Climate classification according to Köppen (1948): 1) Tropical rainforest climate, *Af*; 2) Tropical savanne climate, *As*; 3) Warm summer Mediterranean climate, *Cbs*; and

4) Tropical monsoon climate, *Am* (Chen & Chen, 2013; Köppen, 1884; Peel et al., 2007). For Andean classification (Andes barrier) (0) represents populations in the west side of the Western Andean range and (1) represents populations over or in the east side of this mountain range.

**Supplementary table ST2. DNA microsatellite markers used for genetic structure in *A. cephalotes*.** $N_a$: number of alleles; $H_O$: total observed heterozygosity; $H_E$: total expected heterozygosity; $F_{IS}$: inbreeding coefficient; *HWE*: Hardy-Weinberg equilibrium; *N*: sample size (number of nest). $N_a$ was calculated over all populations. $H_O$ and $H_E$ were calculated as a population mean. Differentiation was estimated by global $F_{ST}$ values per locus and for a multilocus average.

| Locus | $N_a$ | $H_o$ | $H_e$ | $F_{IS}$ | *HWE* | *N* | $F_{ST}$ |
|---|---|---|---|---|---|---|---|
| MAT12 | 20 | 0.91 | 0.87 | -0.06 | NS | 22 | 0.06*** |
| MAT2 | 11 | 0.71 | 0.73 | 0.02 | NS | 15 | 0.12*** |
| MAT10 | 8 | 0.71 | 0.73 | 0.04 | NS | 10 | 0.11*** |
| MAT21 | 8 | 0.60 | 0.59 | -0.02 | NS | 9 | 0.07*** |
| MAT28 | 8 | 0.47 | 0.49 | 0.02 | NS | 9 | 0.19*** |
| MAT4 | 2 | 0.03 | 0.04 | 0.28* | NS | 2 | 0.05*** |
| MAT5 | 4 | 0.25 | 0.24 | -0.03 | NS | 5 | 0.11*** |
| MAT13 | 10 | 0.79 | 0.79 | 0.00 | NS | 14 | 0.07*** |
| MAT20 | 5 | 0.45 | 0.44 | -0.04 | NS | 7 | 0.10*** |
| MAT23 | 8 | 0.64 | 0.60 | -0.07 | NS | 10 | 0.17*** |
| MAT29 | 10 | 0.57 | 0.55 | -0.03 | NS | 12 | 0.19*** |
| MAT15 | 5 | 0.16 | 0.17 | 0.05 | NS | 6 | 0.08*** |
| MAT25 | 11 | 0.76 | 0.78 | 0.03 | NS | 14 | 0.10*** |
| Multilocus | - | 0.54 | 0.54 | -0.01 | NS | - | 0.11* |

Results indicate significance after Bonferroni correction ($\alpha = 0.05$). NS, non-significant; *$P < 0.05$, ***$P < 0.001$.

**Supplementary table ST3. Mitochondrial *COI* neutrality test for sampling locations of *Atta cephalotes* from Pacific, Andean_1 and Andean_2 regions from Colombia.** *D* is Tajima's D and *FS* is Fu's FS. $P_D$ and $P_{FS}$ are the *P*-values for neutrality tests at α = 0.05 and α = 0.02 significance level after Bonferroni correction, respectively. NS: non-significant. N/A: Cannot be calculated (only one allele in the sample).

| | ***D*** | $P_D$ | ***FS*** | $P_{FS}$ |
|---|---|---|---|---|
| *Pacific* | | | | |
| All pops n = 3 | -0.29 | 0.10 | 0.31 | 0.22 |
| BVT | -1.36 | 0.10 | 0.67 | 0.46 |
| QBD | -0.56 | 0.30 | 0.16 | 0.51 |
| GUA | -0.17 | 0.43 | 0.68 | 0.65 |
| *Andean_1* | | | | |
| All pops n = 2 | -0.06 | 0.10 | -0.51 | 0.18 |
| DAG | -0.82 | 0.21 | 0.68 | 0.62 |
| CAL | -0.34 | 0.38 | -0.57 | 0.33 |
| *Andean_2* | | | | |
| All pops n = 5 | -1.06 | 0.10 | -1.83 | 0.09 |
| PPY | 0.32 | 0.69 | 0.15 | 0.42 |
| DOV | -1.72 | 0.15 | -2.68 | 0.02 |
| QBY | -0.40 | 0.28 | 0.13 | 0.28 |
| YOB | 0.00 | 1.00 | 0.00 | N/A |
| MNZ | -0.63 | 0.33 | 0.24 | 0.44 |
| **Total n = 10** | -1.03 | NS | -3.22 | NS |

**Supplementary table ST4.** Comparison of the diversity statistics among regions based on IBE (Pacific, Andean_1 and Andean_2) and IBB (Pacific vs Andean) clustering patterns for both type of markers (microsatellite and *mtCOI*). (*) indicates significance for the comparisons among regions at a $P < 0.05$.

| | IBE | | | | IBB | | |
|---|---|---|---|---|---|---|---|
| **Statistic** | **Pacific** | **Andean_1** | **Andean_2** | ***P*-value** | **Pacific** | **Andean** | ***P*-value** |
| *Nuclear data* | | | | | | | |
| $N_a$ | 6.95 | 4.62 | 4.74 | 0.03* | 6.95 | 4.70 | 0.01* |
| $N_e$ | 4.08 | 2.74 | 2.95 | 0.01* | 4.08 | 2.89 | <0.01* |
| $A_R$ | 6.56 | 4.38 | 4.51 | 0.01* | 6.56 | 4.47 | <0.01* |
| $H_e$ | 0.57 | 0.49 | 0.53 | 0.13 | 0.57 | 0.52 | 0.10 |
| *Mitochondrial data* | | | | | | | |
| $H_d$ | 0.67 | 0.82 | 0.53 | 0.23 | 0.67 | 0.65 | 0.72 |
| π | 0.004 | 0.005 | 0.002 | 0.08 | 0.003 | 0.003 | 0.34 |
| PrivHap | 1 | 0 | 3 | 0.64 | 1 | 3 | 0.86 |

**$N_a$:** mean number of different alleles; **$N_e$:** mean number of effective alleles; **$A_R$:** mean allelic richness; **$H_e$:** mean expected heterozygosity; **$H_d$:** haplotype diversity; **π:** nucleotide diversity; **PrivHap:** number of private haplotypes.

**Supplementary table ST5.** Haplotype (Hap) occurrence across populations.

| | Hap1 | Hap2 | Hap3 | Hap4 | Hap5 | Hap6 | Hap7 | Hap8 | Hap9 | Hap10 | Total |
|---|---|---|---|---|---|---|---|---|---|---|---|
| *Pacific* | | | | | | | | | | | |
| BVT | 8 | 1 | | | | | | | | | 9 |
| QBD | 1 | | 3 | 8 | 1 | | | | | | 13 |
| GUA | 13 | 2 | | 2 | 3 | | | | | | 20 |
| *Andean_1* | | | | | | | | | | | |
| DAG | | | | 2 | 1 | 7 | | | | | 10 |
| CAL | 6 | 5 | | 6 | | | 2 | | | | 19 |
| *Andean_2* | | | | | | | | | | | |
| PPY | 4 | | | 8 | | | 2 | | | | 14 |
| DOV | 16 | | | 1 | | | | 2 | | | 19 |
| QBY | 2 | | | 13 | | | | | | | 15 |
| YOB | 16 | | | | | | | | | | 16 |
| MNZ | 9 | | | | | | | | 1 | 1 | 11 |

**Supplementary table ST6.** Redundancy analysis (RDA) results based on *mtCOI* to determine the relative contribution of environmental, topography and spatial components driving genetic structure in *Atta cephalotes*. Full model (space + temperature + humidity + precipitation + climate + Andean barrier). IBE model (space + temperature + humidity + precipitation + climate). IBB model (space + Andean barrier).

| Model | | | Conditioned model | | |
|---|---|---|---|---|---|
| | F | *P*-value | F | *P*-value | AIC |
| Full (IBD + IBE + IBB) | 0.66 | 0.75 | 0.30 | 0.97 | 2.42 |
| IBE | 0.92 | 0.58 | 0.39 | 0.92 | 1.20 |
| IBB | 1.92 | 0.17 | 0.02 | 0.99 | -1.50 |

* Model selection and its significance is indicated by AIC and *P*-values.

**FIGURES**

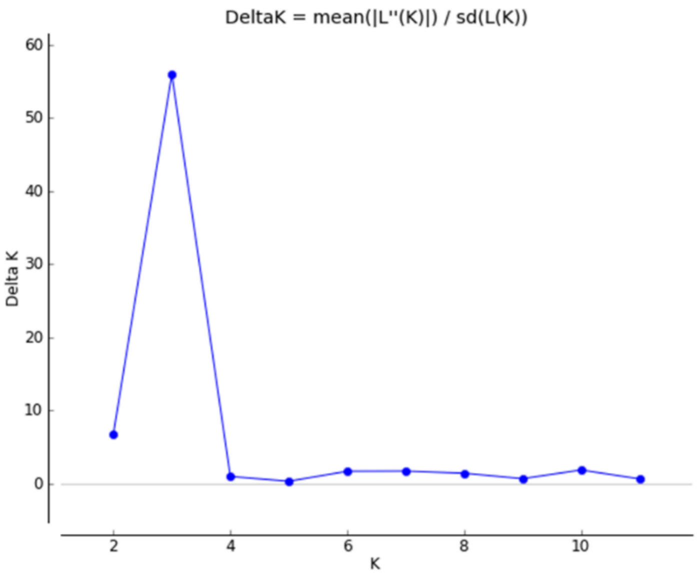


**Supplementary Figure SF1.** Evanno's et al. (2005) plot for detecting the number of the *K* groups that best fit to the data.

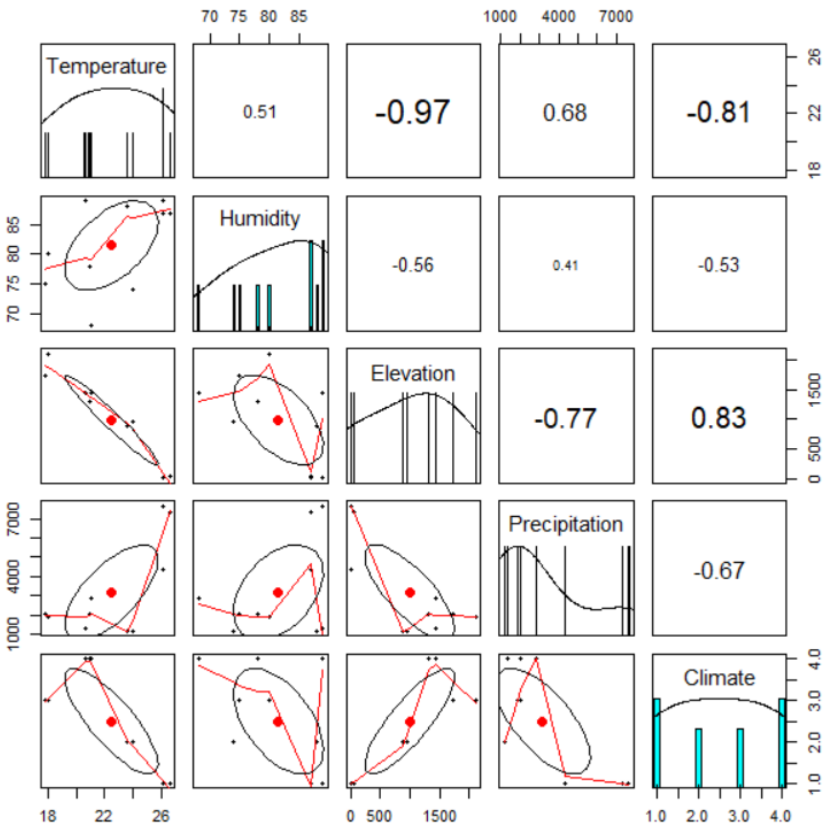


**Supplementary Figure SF2.** Graphic of the correlation test between the environmental variables considered for the RDA analysis. Variables with correlation values > 0.80 were discarded.

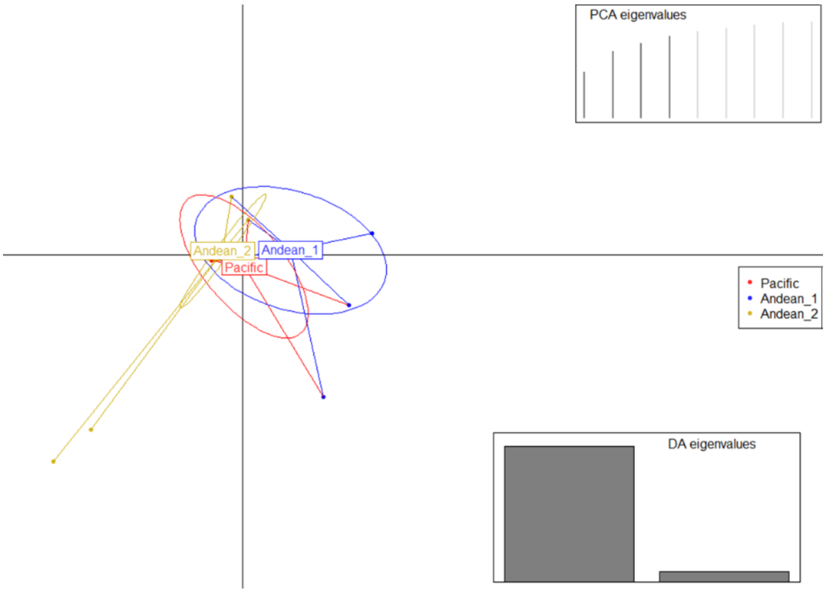


**Supplementary Figure SF3.** DAPC scatterplot showing the genetic structure between *Atta cephalotes* samples, using *mtCOI* data. Key describes the colors attributed to each region and inertia ellipses describe the general distribution of points. Eigenvalues for each PC axis are shown (PC1, vertical; PC2, horizontal). The number of PCA axes retained is shown in the bottom-right inset (gray bars).
